# Transovarial transmission in field caught mosquitoes identifies a mechanism for the establishment of Usutu virus in a temperate country

**DOI:** 10.1101/2024.07.05.602178

**Authors:** Mirjam Schilling, Becki Lawson, Simon Spiro, Madhujot Jagdev, Alexander G.C. Vaux, Robert C. Bruce, Colin J. Johnston, Anthony J. Abbott, Ethan Wrigglesworth, Paul Pearce-Kelly, Andrew A. Cunningham, Jolyon M. Medlock, Nicholas Johnson, Arran J. Folly

## Abstract

Usutu virus (USUV) is an emerging zoonotic flavivirus in Europe, and the first zoonotic mosquito-borne virus to be confirmed in animal hosts in the United Kingdom (UK). Phylogenetic analysis of USUV in the three years following its initial detection in 2020 indicated that the virus is overwintering in the UK. In 2023, USUV was identified outside Greater London for the first time. Therefore, USUV should now be considered endemic within southeast England. Surveillance of avian hosts and mosquito vectors has been insufficient to elucidate the mechanism by which USUV has persisted through temperate winters. It is likely that mosquitoes play a significant role in facilitating the establishment of USUV in temperate areas, as is the case for related mosquito-borne viruses. Here we undertake enhanced targeted vector surveillance at the index site to investigate the role of mosquitoes in facilitating USUV establishment in the UK.

Between 2021 and 2024 inclusive, we detected USUV in host-seeking adult female *Culex pipiens* s.l. (n = 8/554 pools), a key vector of the virus in Europe. During 2023, enhanced surveillance detected transovarial transmission of USUV in wild *Cx. pipiens* s.l. (n = 1/202 pools), by screening adults following captive rearing of mosquito larvae collected from the field. This is, to our knowledge, the first description of vertical transmission of USUV in an arthropod vector. Consequently, transovarial transmission should be considered a viable mechanism for the persistence of USUV in temperate areas. Our results highlight the importance of undertaking detailed vector surveillance, across life stages, to inform the epidemiology of vector-borne viruses.

## Introduction

Global changes in climate are increasing the likelihood of vector-borne disease outbreaks (1, 2). This is especially true for temperate regions that were once considered safe from the establishment of diseases from the tropics or sub-tropics (3). Usutu virus (USUV, family: *Flaviviridae*), first detected in South Africa in 1959 (4), has spread across mainland Europe in recent decades (5) and was first confirmed in the United Kingdom in 2020 (6). The virus was repeatedly confirmed in Greater London in the three years following the initial outbreak, primarily in blackbirds (*Turdus merula*) that were found dead, or euthanised due to morbidity for welfare reasons, and submitted for post-mortem examination. Molecular clock analysis of USUV sequences obtained from infected birds in England across multiple years revealed that they shared a most recent common ancestor, indicating that the virus is overwintering in the UK (7). In 2023, USUV was detected in southeast England outside Greater London for the first time in an infected juvenile blackbird (*Turdus merula*) (8). The recovered genomic sequence was closely related to the Greater London USUV detections, demonstrating previously unrecorded geographic expansion since the 2020 USUV outbreak. Therefore, USUV may now be considered endemic in southeast England.

Typically, USUV infections affect passerines (especially blackbird (*Turdus merula*)) and Strigiformes (8, 9). Indeed, blackbird populations in Greater London appear to have declined by up to 39% since the emergence of USUV (9, 10), indicating that the virus may have had a rapid impact on susceptible hosts. Although human infection is most common asymptomatic, multiple cases of short-term neurological disease have been reported (11). Data from Italy reveal that seropositivity in asymptomatic humans can be as high as 7% (cerebrospinal fluid and serum samples collected between 2008 and 2011 in Modena) (12) or 46% (blood donors in the Lombardy region) (13) in certain areas, compared to 1% in blood donors in the Po river valley in northern Italy reported between 2014 and 2015 (14). These data show that USUV can infect healthy people, with those in areas of higher viral prevalence and where bridge vectors are present, such as forestry workers (18% seroprevalence in northern Italy in 2014-2015) (14) or bird ringers (23.6% seroprevalence in the Netherlands 2021) with increased risk of exposure (15). While there have been no confirmed cases of USUV infection in people in the UK to date, understanding how USUV persists in temperate areas is a key step to inform our understanding of the risk this virus may pose to animal and public health.

As flaviviruses exist in an enzootic cycle, both vector and host likely contribute to the establishment of endemicity in any given area. In order to persist in temperate areas, mosquito-borne viruses must overwinter in either hosts (with the maintenance or recrudescence of virus infection), or vectors to facilitate autochthonous transmission when conditions become permissive. Blackbirds infected with USUV are unlikely to survive the winter (8), while other related species, such as Song Thrush (*Turdus philomelos*), appear to clear the infection with an appropriate antibody response (7). Therefore, it is likely that mosquitoes provide a mechanism for the virus to persist. Both maintenance of infection in diapausing adult mosquitoes and vertical transmission to larvae could be required for overwintering viral persistence in temperate areas. However, there has so far been no detection of virus in wild diapausing mosquitoes or in developing progeny (16).

To improve our understanding of the drivers of USUV establishment, alongside passive surveillance in wild birds (8), we conducted targeted vector surveillance at the site of the initial USUV outbreak in 2020 and where USUV had been detected annually, from 2021 to 2024 inclusive. This comprised 1. Adult mosquito surveillance to improve understanding of the virus circulation and bloodmeal analysis to study transmission networks 2. Overwintering mosquitoes to elucidate mechanisms for virus persistence and 3. Mosquito larvae to investigate potential for transovarial transmission. This approach enabled an increase in virus detection across mosquito life stages, including evidence for vertical transmission, and indicates that *Culex pipiens* s.l. mosquitoes are involved in the persistence of USUV in temperate areas.

## Material and Methods

### Enhanced vector surveillance

To improve our understanding of the contribution of the vector population to the persistence of USUV in the UK enhanced surveillance was conducted. Samples comprised of (1) adult mosquitoes, (2) overwintering mosquitoes and (3) mosquito larvae.

1. Adult female mosquitoes were collected from 2022 and 2023 from ZSL London Zoo, the 2020 index site, using BG-Sentinel 2® traps (Biogents AG, Regensburg, Germany) baited with BG-Lure (Biogents AG) (see Supplementary Table 1). Where possible, captured mosquitoes were collected weekly (2022: Sept; 2023: Jul – Sept) and kept in a -20°C freezer prior to the identification to species level and molecular analysis.
2. Overwintering mosquitoes were collected from November to February at ZSL London Zoo each year, 2021 to 2024 inclusive, in an attempt to elucidate the mechanism of USUV persistence where temperate winters preclude year-round virus transmission (see table 1). A spider catcher was used to collect resting adults. For the last collection in winter 2022/23, and all collections in winter 2023/24, mosquitoes were taken to an insectary at APHA Weybridge and reared for three to four weeks at 25°C and 50% humidity (12:12 light cycle). This was done to end their dormancy and increase their metabolic activity which might reactivate viral replication, facilitating detection using qPCR. Captive mosquitoes were checked daily. Any found dead were recorded and stored at -80°C for later molecular analysis. The first winter collections in 2022 and 2023 did not yield any USUV-positive mosquito pools when the mosquitoes were kept in the laboratory for four weeks. Due to this timeframe resulting in no USUV RNA recorded from mosquitoes in a location shown to have USUV circulation, we hypothesised that mosquitoes might naturally clear the infection over time after diapause. We therefore further adapted our protocol and sampled mosquitoes at 10 days and three weeks post-collection for the mosquitoes captured in December 2023 and January 2024.
3. To identify if USUV transovarial transmission occurs, in 2023 wild mosquito larvae (all first (L1) and second stage (L2) larvae) were collected from June to August 2023 inclusive, at ZSL London Zoo. We targeted standing water pools and additionally established oviposition traps (darkened plastic containers filled two thirds with water) in areas where birds in the zoological collection had seroconverted following autochthonous USUV transmission (7). Trap locations were documented using what3words: (Gravy.Riots.Bugs, Flock.Secret.Files, Formal.Blur.Raves, Common.Swear.Items). Collected larvae were reared to adulthood in an insectary (environmental conditions as above). Adult mosquitoes were checked daily and kept alive for up to four weeks, before being euthanised (by freezing them at -80°C) for molecular analysis. During rearing, any adult mosquitoes found dead were recorded and stored at -80°C prior to later molecular analysis (see Supplementary Table 2).

### RNA extraction and Usutu virus screening

Mosquitoes were homogenised in pools of ≤ ten in 300 µl tissue culture medium using the Qiagen TissueLyser II with 5 mm stainless steel beads (both Qiagen, Manchester, UK) and centrifuged (10,000 rpm/10 min). Total RNA was extracted from 250 µl of the supernatant using TRIzol® (Invitrogen, Life Technologies Limited, Paisley, UK). The precipitated RNA was resuspended in 20 µl nuclease free water and subjected to an USUV-specific RT-qPCR (17). Virus isolation was attempted from homogenate (stored at -80°C) of PCR positive mosquito pools in Vero cells in a BSL3 laboratory at the APHA.

### DNA barcoding

In order to understand potential transmission networks, we attempted to retrieve cytochrome c oxidase I (COX1) sequences of vertebrate hosts from mosquito bloodmeals. Sixteen blood-fed wild mosquitoes caught in September 2022 were processed individually and DNA was extracted from the blood (removed from mosquito using forceps to apply pressure to the abdomen) using the DNeasy Blood & Tissue Kit (Qiagen) according to the manufacturer’s instructions. A 658 bp region located at the 5’ end of the COX1 gene was amplified by PCR as previously established (9, 18). PCR products were visualised on a 1.5% agarose gel and samples of the correct band size were submitted for Sanger sequencing.

### Next generation sequencing and phylogenetic analysis

Mosquito pools that were positive in the USUV-specific RT-qPCR were subjected to next generation sequencing in an attempt to recover a complete virus genome. Sequencing libraries were prepared using the Nextera XT kit (Illumina, Cambridge, UK) and analysed on a NextSeq sequencer (Illumina, Cambridge, UK) with 2 × 150 base paired-end reads. Consensus sequences were generated by using a combination of the Burrows–Wheeler Aligner v0.7.13 and SAMtools v1.9 with a representative USUV genome (MW001216) as a scaffold. Two samples, an adult mosquito pool and an adult reared from a larva, both collected in 2023, produced approximately 5 kb of sequence data that aligned to a USUV reference genome (11 kb -Genbank accession number: MW001216). These sequences clustered with 15 confirmed USUV Africa lineage 3.2 isolates from across mainland Europe using MAFFT v7.471. The resulting alignment was imported into BEAST (v1.10.4) and a Bayesian phylogenetic tree was produced using the GTR+I nucleotide substitution model and 10,000,000 Markov chain Monte Carlo generations. Log files were analysed in Tracer v1.7.1 to check the effective sample size and a 10% burn-in was included (TreeAnnotator v.1.10.4) before being visualised and annotated in FigTree v1.4.4.

## Results

### Enhanced vector surveillance increases USUV detection in mosquitoes

During the mosquito active season in 2022 and 2023 we trapped 156 and 5433 adult mosquitoes, respectively, the majority (92.0%) of which were *Cx. pipiens* s.l. (Figure 1, Supplementary Table 1). Of 55 pools tested from 2022, two pools (3.6%) were positive for USUV by RT-qPCR. In 2023, six of 540 pools (1%) tested positive for USUV by RT-qPCR. The positive pools derived from mosquitoes collected in July and August respectively (7). From one of the 2023 pools we were able to recover a 5088 bp section of the USUV genome for phylogenetic analysis (see below). All pools that tested positive for USUV were *Cx. pipiens* s.l. (Supplementary Table 1). However, we were not able to isolate the virus. Additionally, we identified a total of 16 blood-fed female mosquitoes trapped during September 2022. Blood meal samples from four *Cx. pipiens* amplified a viable avian COXI sequence: southern caracara (*Caracara plancus*) (n = 1), grey-capped emerald dove (*Chalcophas indica*) (n = 1), and scarlet ibis (*Eudocimus ruber*) (n = 2). These are zoological collection birds housed in outdoor enclosures at ZSL London Zoo and are therefore likely to be accessible to host-seeking ornithophagic mosquitoes. No USUV-associated disease has been observed in these collection birds.

**Figure 1:**
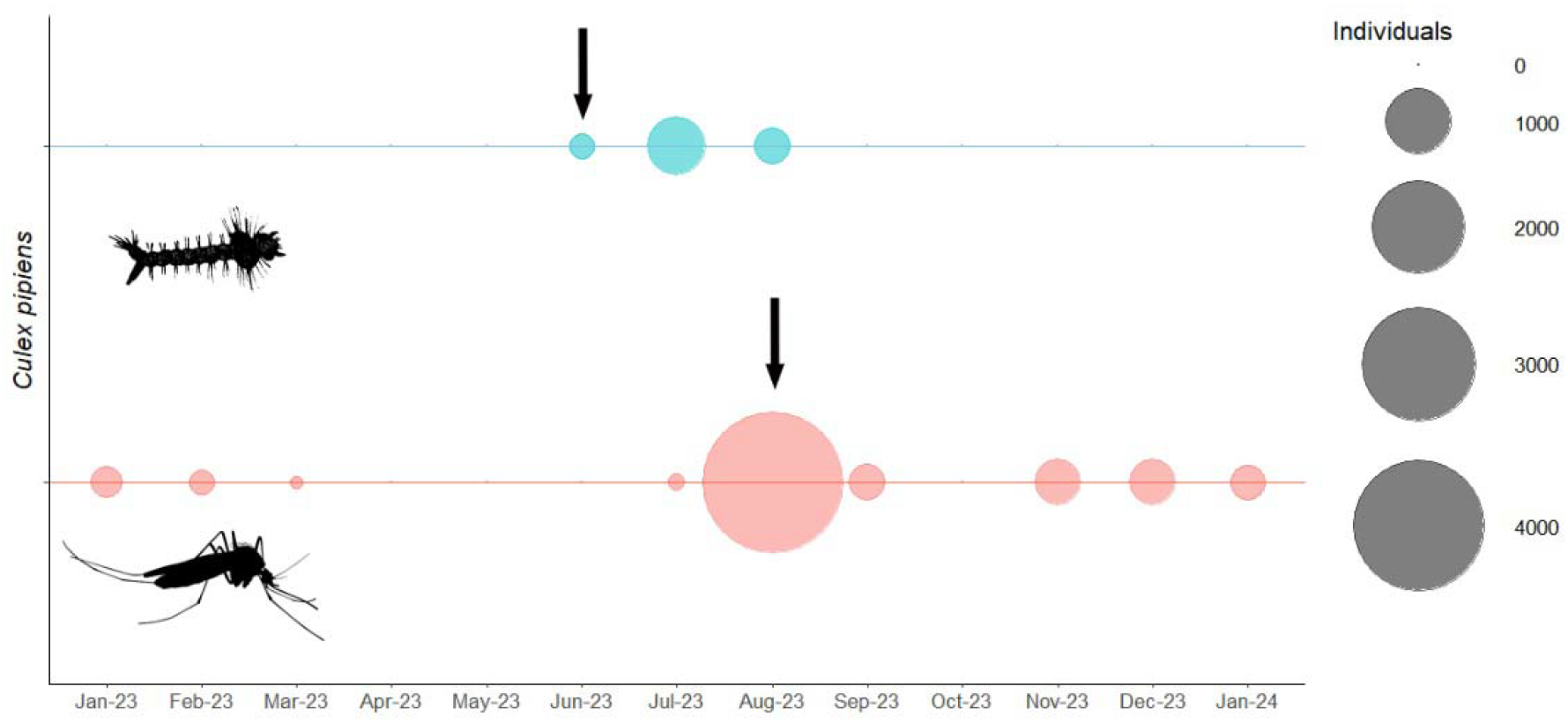
Usutu virus spreads within the *Culex pipiens* s.l. population through transovarial transmission. Bubble plot representing the number of adult *Culex pipiens* (red) and the number of adult *Culex pipiens* reared from larvae collected in the field (blue) between January 2023 and January 2024 inclusive. Arrows mark detections of USUV by RT-qPCR (n=6 adults, n=1 reared from larvae).

### Trialling an adapted protocol to detect USUV in diapausing females

A total of 1950 diapausing females, the majority *Cx. pipiens* s.l. (99.5%), were collected during the winters 2021/22, 2022/23 and 2023/24 (Supplementary Table 1). All of the 40 mosquito pools collected in winter 2021/22, 63 pools collected in winter 2022/23 and 139 collected in winter 2023/24 (total pools n=242 pools) were negative for USUV by RT-qPCR (Supplementary Table 1).

### Transovarial transmission contributes to USUV persistence in Culex pipiens

We reared 1297 adults from wild larvae (L1 and L2) collected in June 2023 (Figure 1, Supplementary Table 2). One pool (n=2 adults) tested positive for USUV (Figure 2), indicating transovarial transmission of the virus. The positive pool contained two male *Cx. pipiens* s.l. that had emerged and died within seven days of field collection from a central location at ZSL London Zoo (what3words location: Gravy.Riots.Bugs). We were unable to isolate virus from homogenate in Vero cells, but next generation sequencing of the positive pool recovered a 5210 bp sequence which mapped to USUV lineage Africa 3.2 (Genbank accession number: MW001216, average read depth = 781). Phylogenetic analysis of a 5088bp region shows that both the transovarially obtained sequence and the adult mosquito sequence (described above) cluster within a USUV lineage Africa 3.2 UK clade (Figure 2).

**Figure 2:**
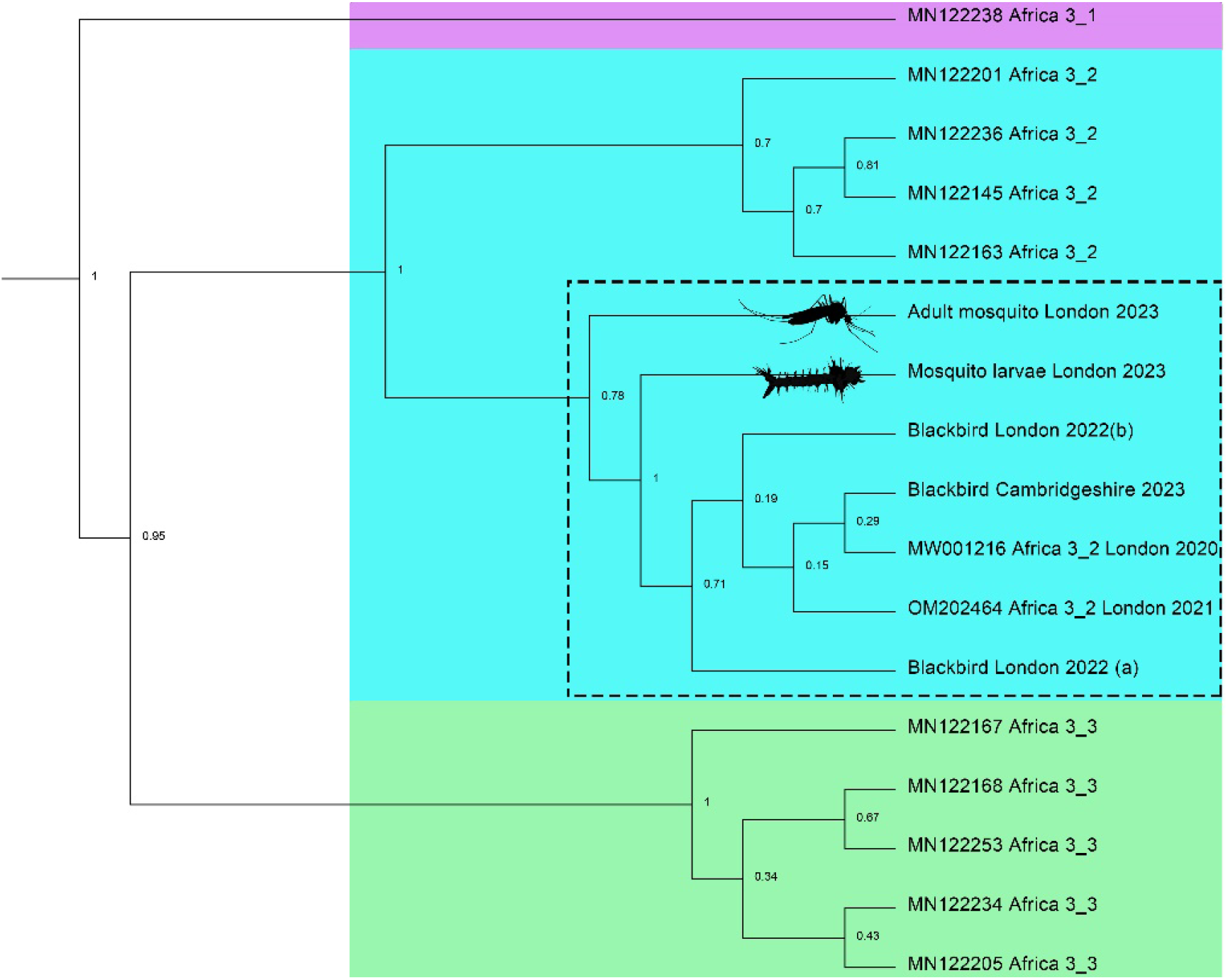
Usutu virus (USUV) sequences from mosquitoes (*Culex pipiens* s.l.) cluster with sequences of isolates from wild birds in England. Bayesian phylogenetic analysis of USUV sequence data (approximately 5kb) retrieved from an adult mosquito pool as well as two adults reared from larva in 2023. The sequences were aligned with 15 USUV Africa lineage isolates from across mainland Europe. Node labels represent posterior probabilities and accession numbers are shown.

## Discussion

In addition to facilitating the detection of USUV RNA in wild-caught adults, our enhanced mosquito surveillance detected USUV RNA in adults which had emerged from wild-caught larvae. This indicates that the virus was transmitted vertically from an infected, gravid, female and suggests transovarial transmission contributes to the establishment of USUV endemicity within vector populations. To our knowledge, this is the first time vertical transmission has been identified for USUV in an arthropod vector.

Even under apparent high virus occurrence at the index site, indicated by the recovery of dead wild passerines infected with USUV and seroconversion of multiple birds in the zoological collection (7), our study found the prevalence of USUV detectable by RT-qPCR was extremely low in the mosquito community. This is in line with previous results from earlier mosquito surveillance at the index site in 2021 (7), and similar studies performed in mainland Europe where USUV is circulating (7, 19). It is, however, unclear if these results are caused by a genuine low prevalence of USUV in the vector population, or if virus is maintained at titres within the insect host below the level of detection of our RT-qPCR assay. Studies on related flaviviruses using next generation sequencing technologies report difficulty in detecting viral RNA without targeted amplification of flavivirus RNA prior to sequencing, even in areas with known high virus prevalence, suggesting that virus titres in vectors are naturally low (20). Our investigations of USUV, which targeted the different stages (collected as larvae and adults, and tested as adults) of the mosquito life cycle help to elucidate the contributions of arthropod vectors to the persistence and endemicity of USUV in temperate areas.

We detected USUV RNA in two pools of adult mosquitoes collected in September 2022 and six pools in August 2023, the period of peak abundance of adult mosquitoes in Greater London (7, 21). Combining vector abundance data with USUV prevalence will help to identify areas and times of the year when there is an increased risk of virus exposure, to raise awareness amongst medical and veterinary communities and target mitigation measures to safeguard health, such as deploying targeted mosquito control. Additionally, our blood meal analyses show that *Cx. pipiens* is an opportunistic, ornithophagic mosquito, meaning that a diverse range of birds may inadvertently be exposed to virus. This is in line with confirmed USUV detections in the UK and across mainland Europe as well as seroconversion following autochthonous USUV transmission, identified in a range of birds held in the zoological collection at ZSL London Zoo (7, 9). Consequently, it is likely that species of birds in addition to wild passerines may be contributing to the enzootic cycle in a given area.

While USUV is emerging across Europe and is considered endemic in many countries, detection of overwintering flaviviruses in diapausing mosquitoes is rarely recorded (16, 22). Since mRNA expression during diapause is reduced and a transcriptional program leads to metabolic repression (23, 24), it is likely that viral RNA is maintained at low titres in mosquito cells until conditions for replication become favourable. Therefore, our methodology opted to rear overwintering females in the laboratory under optimum summer conditions for up to four weeks after field sampling. However, these conditions did not result in detection of USUV RNA, even after readjusting the protocol with a group of females being tested as early as 10 days after collection. Therefore, it is likely that either there was no USUV RNA or no active infection in the mosquito community we sampled, or that effective USUV replication would have required longer than our protocol allowed (25, 26). Future studies might also attempt to bloodfeed mosquitoes after diapause to investigate whether this initiates virus replication. If successful, this might imply that in the field viruses emerge in the first generation of larvae rather than in the overwintering mosquito. Infection prevalence for the overwintering of a related flavivirus has been published to be as low as 0.016% and might be similar for USUV (27). Our recommendation is that laboratory-based experiments investigating the ability of arboviruses to persist through, and replicate after, diapause would provide a deeper understanding of USUV vector infection dynamics. However, we note that our overwintering survey was conducted in a restricted geographic area in central London and more mosquito hibernacula in a wider area should be investigated.

There are likely multiple drivers for the establishment of mosquito-borne pathogens in new areas. Transovarial transmission in arthropod vectors, generally observed at relatively low levels in field samples, has been implicated as a potential persistence mechanism for flaviviruses during periods of unsuitable climatic conditions or in the absence of vertebrate hosts (28, 29). Through our enhanced surveillance, we detected USUV in adult mosquitoes that had emerged from field caught larvae. To our knowledge, this is the first time vertical transmission has been reported for USUV in arthropod vectors. As vertical transmission allows virus spread within the mosquito population independent of any host activity, this may be a key mechanism for the establishment of this virus in temperate areas. One could also speculate that our detection of transovarial transmission of USUV in a larval pool in June is indirect indication for overwintering, as these might have been the first larvae that developed from eggs laid after diapause (21).

As some native UK mosquito species are competent vectors for African USUV lineages (30) our results emphasize the need to better understand the enzootic cycle of USUV in the UK, with a particular focus on the involvement of mosquitoes in viral persistence. This is especially true if bridge vectors are involved in enzootic cycles as this could have negative implications for public health. Our data implicate both adult mosquitoes and developing larvae in the establishment of endemicity in temperate areas. Since flaviviruses are set to continue to spread across Europe in the coming years (31), our study highlights the value of enhancing vector surveillance to help inform disease ecology and epidemiology and to identify areas that warrant targeted mosquito control to minimise infection risk for humans and animals.

## Author contributions

### Funding

This study was supported by grant SV3045 from Department for Environment, Food & Rural Affairs (Defra) and the Devolved Administrations of Scotland and Wales and a UKRI funded project Vector-Borne RADAR (BB/X017990/1). Financial support for the Garden Wildlife Health project comes in part from the Defra, the Welsh Government and the APHA Diseases of Wildlife Scheme Scanning Surveillance Programme (Project ED1058), and from the Garfield Weston Foundation and the Universities Federation for Animal Welfare. BL and AAC receive financial support from Research England.

## Acknowledgments

The authors would like to thank Fiona McCracken for assisting with RNA extractions and the central sequencing unit (Animal and Plant Health Agency).

## Authors’ contributions

Manuscript first draft: MS, AJF; Investigations: MS, BL, SS, MJ, RCB, CJJ, AJA, AAC, JMM, AJF; Technical and methodological support: MJ, AGJV, RCB, CJJ, AJA, EW, PPK, NJ; Study design and analysis: MS, AJF; Project management / supervision: AJF, MS, JMM; Funding: AJF, NJ, BL, JMM; Manuscript review and edit: all authors.

## Data availability

### Ethics statement

The authors confirm that the ethical policies of the journal, as noted on the journal’s author guidelines page, have been adhered to. Passive surveillance was conducted through post-mortem examination of wild birds found dead, or euthanised for welfare reasons.

## Supplementary Tables

**Supplementary Table 1:** Adult mosquitoes collected at the index site in Greater London, UK, October 2021 to January 2024 inclusive.

| Year | Month | *Culex pipiens* trapped | Pools | USUV positive pools | *Culiseta annulata* trapped | Pools | USUV positive pools |
| --- | --- | --- | --- | --- | --- | --- | --- |
| 2021 | October | 30 | 3 | 0 | 0 | 0 | 0 |
| 2021 | November | 40 | 4 | 0 | 0 | 0 | 0 |
| 2021 | December | 12 | 3 | 0 | 2 | 2 | 0 |
| 2022 | January | 171 | 26 | 0 | 2 | 2 | 0 |
| 2022 | September | 151 | 51 | 2 | 5 | 4 | 0 |
| 2022 | October | 2 | 2 | 0 | 1 | 1 | 0 |
| 2022 | November | 32 | 6 | 0 | 1 | 1 | 0 |
| 2022 | December | 43 | 7 | 0 | 0 | 0 | 0 |
| 2023 | January | 221 | 25 | 0 | 0 | 0 | 0 |
| 2023 | February | 143 | 21 | 0 | 0 | 0 | 0 |
| 2023 | March | 32 | 4 | 0 | 0 | 0 | 0 |
| 2023 | July | 61 | 11 | 0 | 0 | 0 | 0 |
| 2023 | August | 4600 | 460 | 6 | 390 | 39 | 0 |
| 2023 | September | 300 | 32 | 0 | 50 | 5 | 0 |
| 2024 | November | 482 | 54 | 0 | 3 | 3 | 0 |
| 2023 | December | 480 | 51 | 0 | 1 | 1 | 0 |
| 2024 | January | 278 | 29 | 0 | 1 | 1 | 0 |

**Supplementary Table 2:** Adult mosquitoes reared from larvae collected at the index site in Greater London, UK, in 2023.

| Month | Adult *Culex pipiens* emerged | Pools | USUV positive pools | Adult *Culiseta annulata* emerged | Pools | USUV positive pools |
| --- | --- | --- | --- | --- | --- | --- |
| June | 137 | 53 | 1 | 2 | 1 | 0 |
| July | 789 | 83 | 0 | 29 | 13 | 0 |
| August | 304 | 66 | 0 | 36 | 14 | 0 |

## References

1. Paz S. Climate change impacts on vector-borne diseases in Europe: Risks, predictions and actions. The Lancet Regional Health – Europe. 2021;1.

2. de Souza WM, Weaver SC. Effects of climate change and human activities on vector-borne diseases. Nature Reviews Microbiology. 2024.

3. Paz S. Climate change: A driver of increasing vector-borne disease transmission in non-endemic areas. PLOS Medicine. 2024;21(4):e1004382.

4. Williams MC, Simpson DI, Haddow AJ, Knight EM. THE ISOLATION OF WEST NILE VIRUS FROM MAN AND OF USUTU VIRUS FROM THE BIRD-BITING MOSQUITO MANSONIA AURITES (THEOBALD) IN THE ENTEBBE AREA OF UGANDA. Ann Trop Med Parasitol. 1964;58:367–74.

5. Siljic M, Sehovic R, Jankovic M, Stamenkovic G, Loncar A, Todorovic M, et al. Evolutionary dynamics of Usutu virus: Worldwide dispersal patterns and transmission dynamics in Europe. Frontiers in Microbiology. 2023;14.

6. Folly AJ, Lawson B, Lean FZ, McCracken F, Spiro S, John SK, et al. Detection of Usutu virus infection in wild birds in the United Kingdom, 2020. Euro Surveill. 2020;25(41).

7. Folly AJ, Sewgobind S, Hernández-Triana LM, Mansfield KL, Lean FZX, Lawson B, et al. Evidence for overwintering and autochthonous transmission of Usutu virus to wild birds following its redetection in the United Kingdom. Transbound Emerg Dis. 2022.

8. Giglia G, Agliani G, Munnink BBO, Sikkema RS, Mandara MT, Lepri E, et al. Pathology and Pathogenesis of Eurasian Blackbirds (Turdus merula) Naturally Infected with Usutu Virus. Viruses. 2021;13(8):1481.

9. Lawson B, Robinson RA, Briscoe AG, Cunningham AA, Fooks AR, Heaver JP, et al. Combining host and vector data informs emergence and potential impact of an Usutu virus outbreak in UK wild birds. Sci Rep. 2022;12(1):10298.

10. BTO. Population trend graphs [Available from: https://www.bto.org/our-science/projects/breeding-bird-survey/latest-results/population-trend-graphs.

11. Cadar D, Simonin Y. Human Usutu Virus Infections in Europe: A New Risk on Horizon? Viruses. 2022;15(1).

12. Grottola A, Marcacci M, Tagliazucchi S, Gennari W, Di Gennaro A, Orsini M, et al. Usutu virus infections in humans: a retrospective analysis in the municipality of Modena, Italy. Clin Microbiol Infect. 2017;23(1):33–7.

13. Percivalle E, Cassaniti I, Sarasini A, Rovida F, Adzasehoun KMG, Colombini I, et al. West Nile or Usutu Virus? A Three-Year Follow-Up of Humoral and Cellular Response in a Group of Asymptomatic Blood Donors. Viruses. 2020;12(2).

14. Percivalle E, Sassera D, Rovida F, Isernia P, Fabbi M, Baldanti F, et al. Usutu Virus Antibodies in Blood Donors and Healthy Forestry Workers in the Lombardy Region, Northern Italy. Vector Borne Zoonotic Dis. 2017;17(9):658–61.

15. de Bellegarde de Saint Lary C, Kasbergen LMR, Bruijning-Verhagen P, van der Jeugd H, Chandler F, Hogema BM, et al. Assessing West Nile virus (WNV) and Usutu virus (USUV) exposure in bird ringers in the Netherlands: a high-risk group for WNV and USUV infection? One Health. 2023;16:100533.

16. Blom R, Schrama MJJ, Spitzen J, Weller BFM, van der Linden A, Sikkema RS, et al. Arbovirus persistence in North-Western Europe: Are mosquitoes the only overwintering pathway? One Health. 2023;16:100467.

17. Jöst H, Bialonski A, Maus D, Sambri V, Eiden M, Groschup MH, et al. Isolation of usutu virus in Germany. Am J Trop Med Hyg. 2011;85(3):551–3.

18. Hernández-Triana LM, Brugman VA, Prosser SWJ, Weland C, Nikolova N, Thorne L, et al. Molecular approaches for blood meal analysis and species identification of mosquitoes (Insecta: Diptera: Culicidae) in rural locations in southern England, United Kingdom. Zootaxa. 2017;4250(1):67–76.

19. Rau J, Köchling K, Schäfer M, Tews BA, Wylezich C, Schaub GA, et al. Viral RNA in Mosquitoes (Diptera: Culicidae) Collected between 2019 and 2021 in Germany. Viruses. 2023;15(12).

20. Thannesberger J, Rascovan N, Eisenmann A, Klymiuk I, Zittra C, Fuehrer HP, et al. Viral metagenomics reveals the presence of novel Zika virus variants in Aedes mosquitoes from Barbados. Parasit Vectors. 2021;14(1):343.

21. Seechurn N, Herdman MT, Hernandez-Colina A, Vaux AGC, Johnston C, Berrell M, et al. Field-based assessments of the seasonality of Culex pipiens sensu lato in England: an important enzootic vector of Usutu and West Nile viruses. Parasites & Vectors. 2024;17(1):61.

22. Rudolf I, Betášová L, Blažejová H, Venclíková K, Straková P, Šebesta O, et al. West Nile virus in overwintering mosquitoes, central Europe. Parasit Vectors. 2017;10(1):452.

23. Denlinger DL. REGULATION OF DIAPAUSE. Annual review of entomology. 2002;47(1):93–122.

24. Sim C, Kang DS, Kim S, Bai X, Denlinger DL. Identification of FOXO targets that generate diverse features of the diapause phenotype in the mosquito Culex pipiens. Proc Natl Acad Sci U S A. 2015;112(12):3811–6.

25. Ferreira PG, Tesla B, Horácio ECA, Nahum LA, Brindley MA, de Oliveira Mendes TA, et al. Temperature Dramatically Shapes Mosquito Gene Expression With Consequences for Mosquito-Zika Virus Interactions. Front Microbiol. 2020;11:901.

26. Ciota AT, Keyel AC. The Role of Temperature in Transmission of Zoonotic Arboviruses. Viruses. 2019;11(11).

27. Kampen H, Tews BA, Werner D. First Evidence of West Nile Virus Overwintering in Mosquitoes in Germany. Viruses. 2021;13(12).

28. Janjoter S, Kataria D, Yadav M, Dahiya N, Sehrawat N. Transovarial transmission of mosquito-borne viruses: a systematic review. Frontiers in Cellular and Infection Microbiology. 2024;13.

29. Lequime S, Paul RE, Lambrechts L. Determinants of Arbovirus Vertical Transmission in Mosquitoes. PLoS Pathog. 2016;12(5):e1005548.

30. Hernández-Triana LM, de Marco MF, Mansfield KL, Thorne L, Lumley S, Marston D, et al. Assessment of vector competence of UK mosquitoes for Usutu virus of African origin. Parasit Vectors. 2018;11(1):381.

31. Vilibic-Cavlek T, Petrovic T, Savic V, Barbic L, Tabain I, Stevanovic V, et al. Epidemiology of Usutu Virus: The European Scenario. Pathogens. 2020;9(9).

